# Hypereosinophilia causes progressive cardiac pathologies in mice

**DOI:** 10.1101/2022.05.04.490445

**Authors:** Nicola Laura Diny, Megan Kay Wood, Taejoon Won, Monica Vladut Talor, Clarisse Lukban, Djahida Bedja, Nadan Wang, C. Conover Talbot, Brian Leei Lin, Daniela Čiháková

## Abstract

Hypereosinophilic syndrome is a progressive disease with extensive eosinophilia that results in organ damage. Cardiac pathologies are the main reason for its high mortality rate. A better understanding of the mechanisms of eosinophil-mediated tissue damage would benefit therapeutic development. Here, we describe the cardiac pathologies that developed in a mouse model of hypereosinophilic syndrome. These IL-5 transgenic mice exhibited decreased left ventricular function at a young age which worsened with age. Mechanistically, we demonstrated infiltration of activated eosinophils into the heart tissue that led to an inflammatory environment. Gene expression signatures showed tissue damage as well as repair and remodeling processes. Cardiomyocytes from IL-5Tg mice exhibited significantly reduced contractility relative to WT controls. This impairment may result from the inflammatory stress experienced by the cardiomyocytes and suggest that dysregulation of contractility and Ca^2+^ reuptake in cardiomyocytes contributes to cardiac dysfunction at the whole organ level in hypereosinophilic mice.

**Teaser:** Too many eosinophils cause inflammation in the heart and change cardiomyocyte contraction leading to poor heart function.

## Introduction

Hypereosinophilic syndromes (HES) are a range of diseases marked by persistent eosinophilia in the absence of primary causes (such as allergies or parasitic infection) (*1-4*). Tissue infiltration of eosinophils can damage multiple organ systems, including the lung, skin, and gastrointestinal tract. Involvement of the cardiovascular system occurs in up to 50% of HES patients and is a major contributor to morbidity and mortality (*5*). The characteristic presentation is a progression from eosinophilic endomyocarditis that damages the endothelial lining to mural thrombi followed by endomyocardial fibrosis and valvular disease (*6, 7*). Current targeted immunotherapies against eosinophils (anti-IL-5 or anti-IL-5 receptor) have improved patient outcomes and reduced the need for non-specific immunosuppressive or cytotoxic therapies and thereby reduced treatment-related comorbidities (*8, 9*). They also clearly demonstrate the causal role of eosinophils in this disease. Nevertheless, current therapies do not eliminate morbidity and there is an ongoing need for the development of preclinical models and mechanistic understanding of the molecular pathology (*10, 11*).

Eosinophils are multi-potent effector cells that can modulate immune responses and tissue remodeling but also cause tissue damage. Their granules contain cytotoxic proteins that can be released by degranulation and damage cells directly, by rupturing membranes or through the production of reactive oxygen species (*12*). Eosinophils also produce many different cytokines, chemokines, and lipid mediators that can recruit and activate other immune cells (*13*). How eosinophils damage the heart in hypereosinophilic syndromes is not fully understood. Proposed mechanisms include direct tissue damage by the granule proteins major basic protein (MBP) and eosinophil cationic protein (ECP) that can be found deposited in areas of eosinophil infiltration in the endocard and myocard (*14*). Eosinophils may also promote coagulation on the ventricular endothelial lining, in part by impairing the function of thrombomodulin (*7, 15*). More recently, the potential of eosinophils to promote the formation of thrombi and its association with cardiovascular disease has been clearly demonstrated. Their capacity to generate thrombin is based on the enzymatic generation of a procoagulant phospholipid surface (*16*). Eosinophils may also contribute to the fibrotic end stage of HES-mediated heart disease. An association between eosinophils and fibrosis is widely recognized across different diseases (*17-19*) and multiple mechanisms are likely at play. Eosinophils express transforming growth factor β and can promote proteoglycan accumulation, extracellular matrix synthesis, and fibroblast proliferation (*20-24*). Which of these processes is important in HES is not known.

To generate a model of hypereosinophilia in mice, IL-5 is overexpressed under the CD3δ promoter (*25*). This leads to the development of massive eosinophilia in the blood and eosinophil infiltration in most organs. The first description of these IL-5Tg mice describes frequent premature death, with 70% dying by 12 months of age (*25*). Cardiomegaly and eosinophil infiltration of the heart was noted in some of the dead mice but not further analyzed. Here, we characterize the cardiac phenotype in detail, analyzing mice from 6 up to 40 weeks of age. Cardiac pathologies worsened with age and included left ventricular dysfunction, increased heart weight, abnormalities of the conductive system and immune infiltration that resulted in tissue damage and remodeling.

## Results

### Hypereosinophilic mice have impaired cardiac function

Given the frequent cardiac complications of patients with hypereosinophilic syndrome, we sought to study the effects of hypereosinophilia on the heart in mice. We used mice with transgenic expression of Il5 (IL-5Tg) which develop hypereosinophilia in blood and multiple organs that worsens with age (*25*). Already in young adult hypereosinophilic mice impaired cardiac function was evident by echocardiography (Figure 1). At 6-9 weeks of age, IL-5Tg mice had increased left ventricular end diastolic diameter (LVEDD) and end-systolic diameter (LVESD), which resulted in decreased ejection fraction (EF) (Figure 1E-G). There was no significant difference in the heart rate (Figure 1H). Hypereosinophilic mice also had increased left ventricular wall thickness of the interventricular septum (IVSD) and posterior wall (PWTED, Figure 1I, J). However, the relative wall thickness was similar in WT and IL-5Tg mice (Figure 1K). As a result of the increased ventricular diameters and wall thickness, the LV mass calculated from the echocardiography analysis was also increased in IL-5Tg mice (Figure 1L), which was not a consequence of higher body weight (Figure 1M). On histological examination, IL-5Tg hearts showed no large-scale alterations but eosinophils were visible in the tissue (Figure 1 O,N). These findings suggest a connection between reduced cardiac function and eosinophils in the heart.

**Figure 1.**
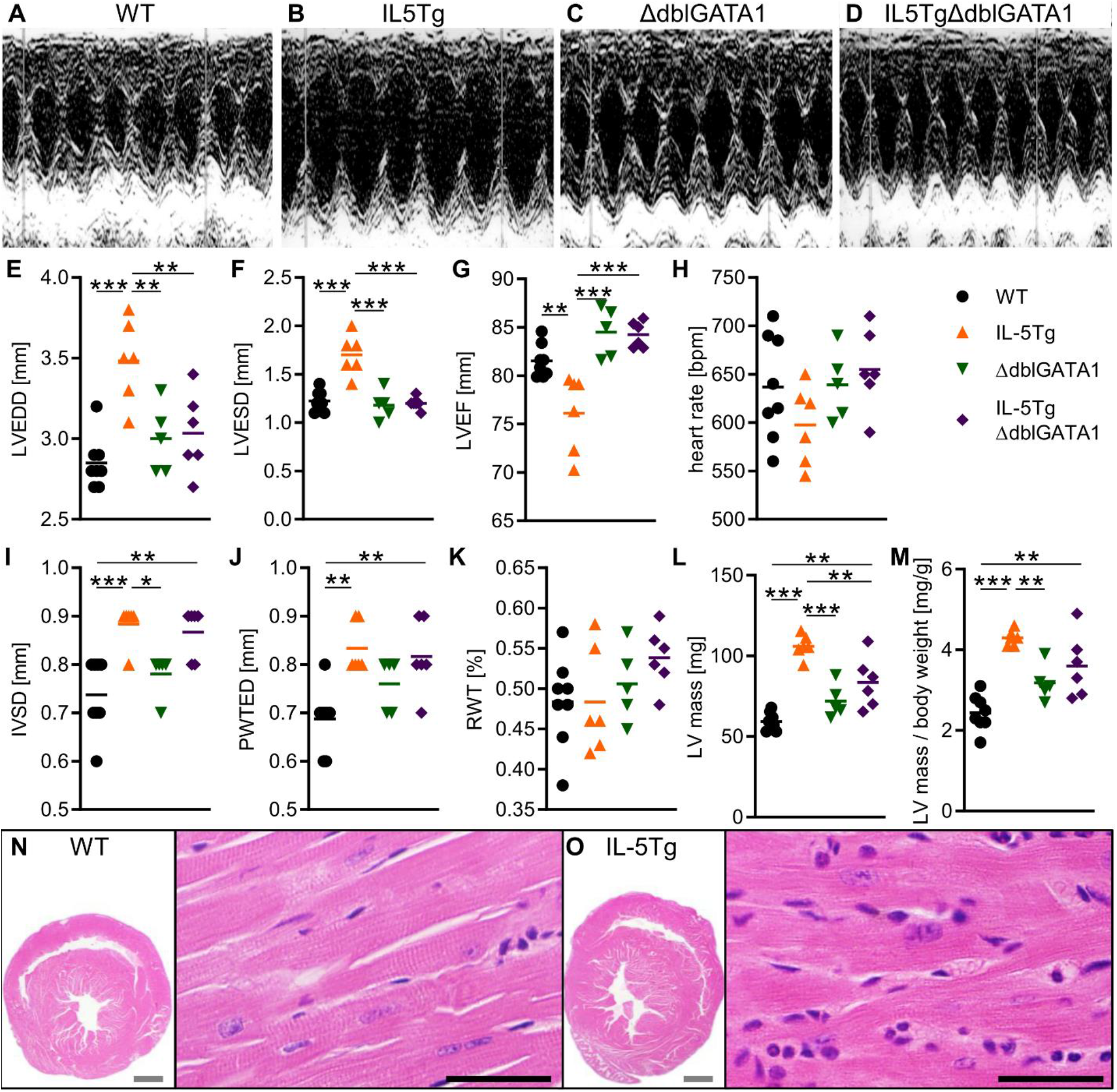
Young naïve hypereosinophilic mice have decreased cardiac function. A-M) Echocardiography was performed on 6-9 week old naïve mice of the indicated genotypes. A-D) representative M-mode images. E-M) Data are representative of 3 independent experiments with 5-9 mice/ group and were analyzed by one-way ANOVA with Tukey post test comparing all groups against each other. N-O) Representative images of H&E-stained heart sections. Grey scale bars: 1mm, black scale bars: 40µm.

IL-5Tg mice have an expansion of T cells and B cells in addition to hypereosinophilia (*25*). IL-5 may also have unknown effects on the heart that are independent from eosinophils. To differentiate between the effects of IL-5 and eosinophils, we analyzed cardiac function in IL-5Tg eosinophil-deficient mice (IL-5TgΔdblGATA1) and eosinophil-deficient controls (ΔdblGATA1). Echocardiography readouts such as LV diameters and EF were similar in IL-5TgΔdblGATA1, ΔdblGATA1, and WT mice and significantly higher in IL-5Tg mice compared to all other groups (Figure 1E-M), confirming that eosinophils, rather than IL-5, were responsible for the observed phenotype. These data show that hypereosinophilia is associated with impaired cardiac function already in young adult mice.

### Cardiac dysfunction in hypereosinophilic mice worsens with age

We hypothesized that the effects of hypereosinophilia on cardiac function may worsen over time. Indeed, the heart weight / body weight ratio increased dramatically with age in IL-5Tg, but not WT mice (Figure 2A). Linear regression of age with heart weight / body weight ratio was not significant for WT mice (r^2^=0.0001, p=0.9344) but highly significant for IL-5Tg mice (r^2^=0.3650, p<0.0001). This increase in relative heart weight likely reflects deteriorating cardiac pathology as the mice age. In HES patients, left ventricular hypertrophy is a common finding (*7*). This prompted us to investigate whether cardiomyocytes showed signs of hypertrophy. We analyzed cardiomyocyte size using wheat germ agglutinin assay. However, no difference in average cardiomyocyte diameter was observed between the different genotypes (Figure 2B-D). We therefore assessed cardiac function by echocardiography and electrocardiography in older mice (age 20-30 weeks). Older hypereosinophilic mice showed increased LV end-systolic and end-diastolic diameters and decreased ejection fraction compared to WT mice (Figure 2E-G). Moreover, the ejection fraction was below 50% for several older hypereosinophilic mice, which we had not observed in young mice (Figure 1G). On average, the LV wall thickness was similar between WT and IL-5Tg mice, some IL-5Tg mice exhibited particularly thick or particularly thin LV intraventricular septum and LV posterior walls (Figure 2H-J). This suggests that different cardiac pathologies may have developed in these mice. In younger hypereosinophilic mice we had observed an insignificant trend towards a slower heart rate in the IL-5Tg mice. In older mice, the heart rate was significantly lower in hypereosinophilic mice (Figure 2K). Similar to young hypereosinophilic mice, the LV mass was increased in older IL-5Tg compared to WT mice (Figure 2L, M). To prove that cardiac pathology worsened with age in IL-5Tg mice, we conducted serial echocardiography in a cohort of mice. Indeed, LV diameters and ejection fraction worsened already from 8 to 16 weeks of age in IL-5Tg mice and LV mass increased (Supplemental Figure 1). In conclusion, left ventricular pathology in IL-5Tg mice worsens with age.

**Figure 2.**
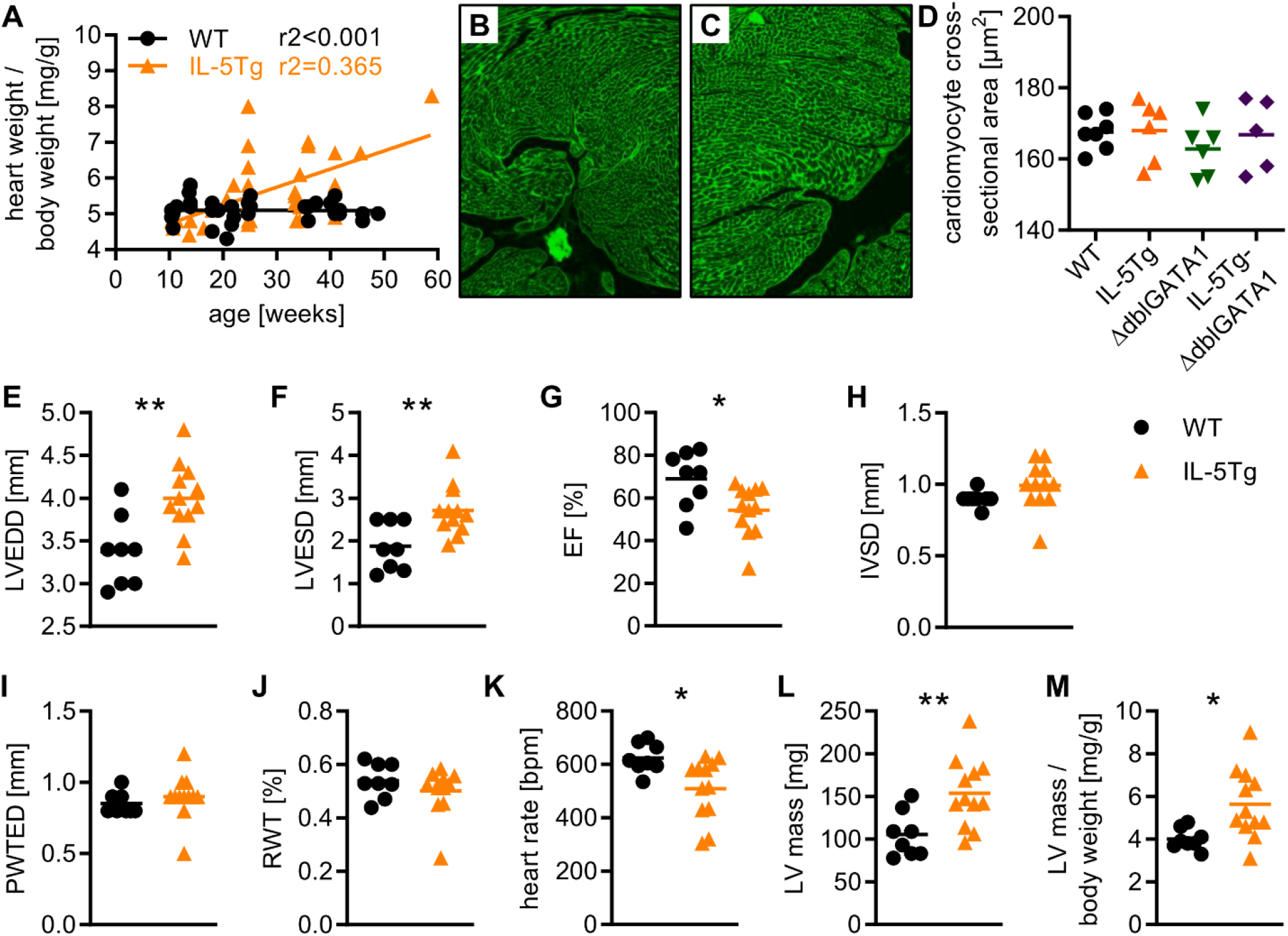
Age exacerbates cardiac pathology in hypereosinophilic mice. A) Heart weight / body weight ratio in relation to age. Data are from 73 mice pooled from multiple experiments and were analyzed by simple linear regression. The slopes for WT compared to IL-5Tg mice are significantly different (p<0.0001). B, C) Representative images of B) WT and C) IL5Tg mice showing wheat germ agglutinin staining of transverse cardiac sections. D) Quantification of average cardiomyocyte diameter in the indicated genotypes. Data are from one experiment with 5-7 mice/group (age 9-14 weeks). E-M) Echocardiography was performed on mice of 20-30 weeks of age. Data from 8-12 mice per genotype were analyzed by t-test.

### Hypereosinophilic mice develop infiltration of the lungs but not pulmonary hypertension

When examining an old IL-5Tg mouse, we noticed enlargement of the right ventricle. The dilation caused the interventricular septum to push into the left ventricle at the end of systole (Supplemental Figure 2A-D, Supplemental Movie 1, 2). This was suggestive of right heart enlargement secondary to pulmonary resistance. We therefore investigated the lungs, which showed diffuse eosinophilic inflammation (Supplemental Figure 2E). This finding prompted us to analyze lung inflammation in a larger cohort of mice (age 10-41 weeks) to determine its frequency among hypereosinophilic mice. While WT mice showed no or only mild lung inflammation, almost all IL-5Tg mice had varying degrees of inflammatory infiltrate (Figure 3A-E). Pulmonary infiltration increased with age in IL5Tg mice (Figure 3E). To characterize the infiltrating cells, we analyzed the lungs by flow cytometry. In 30-41-weeks-old mice, lung weight was increased in IL-5Tg mice (Figure 3F). We therefore compared infiltrating immune cell numbers per mg tissue.

**Figure 3.**
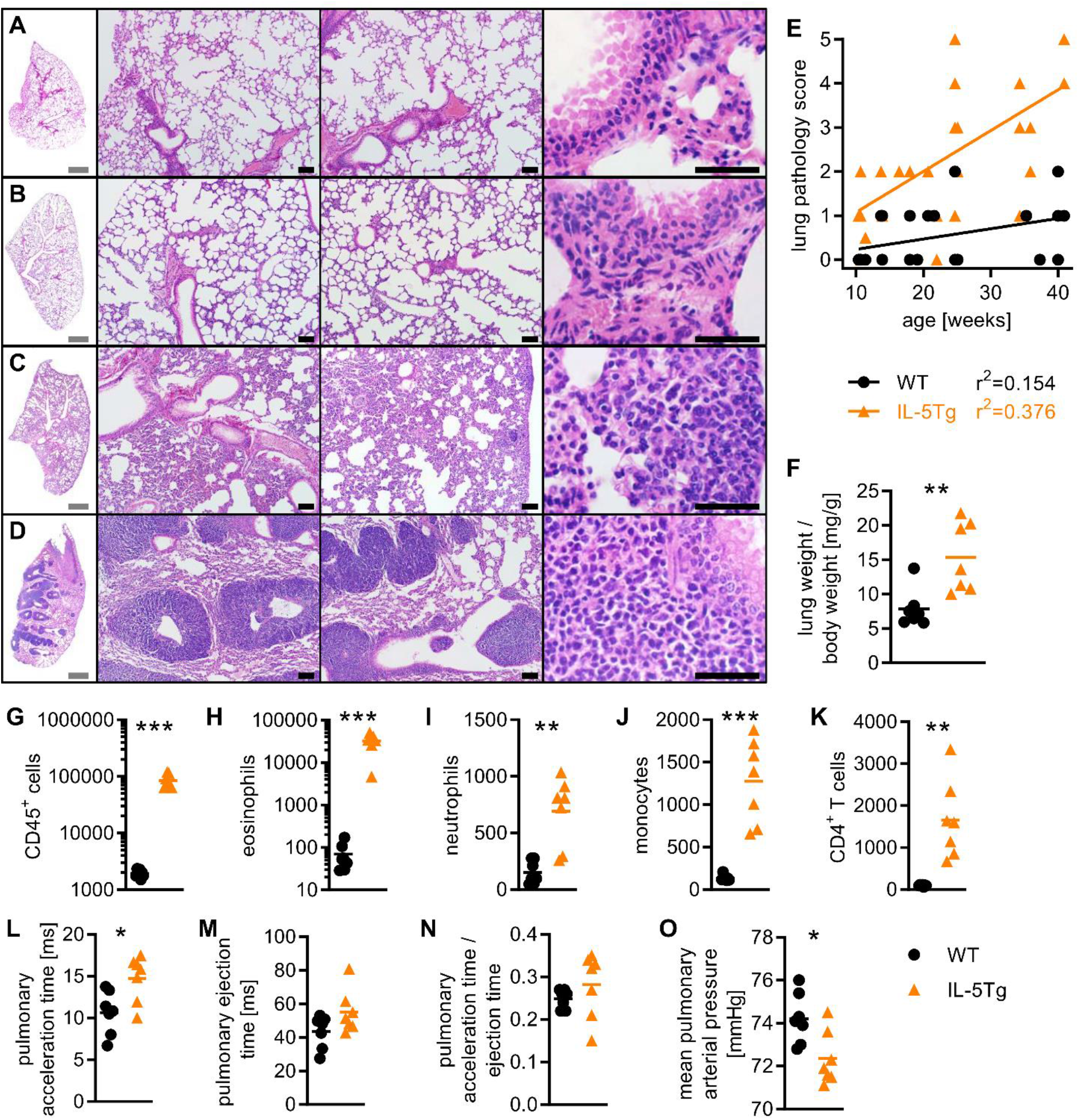
Pulmonary inflammation is common in hypereosinophilic mice but does not lead to pulmonary hypertension. A) WT mice and B-D) IL5Tg mice with different lung pathology scores. A) score 0, B) score 1, C) score 3, D) sore 5. Grey scale bars: 1mm, black scale bars: 40µm. E) Lung pathology score in relation to age. Data are from 57 mice pooled from 2 independent experiments and were analyzed by simple linear regression. The slopes were significantly different for WT compared to IL-5Tg mice (p=0.0072). F-K) Lungs of 30-41-weeks-old mice were analyzed. G-K) Infiltrating immune cells were quantified by flow cytometry and normalized to tissue weight (cells/ mg tissue). L-O) Echocardiography of mice aged 30-41 weeks. F-O) Data are from 7 mice per group and were analyzed by t-test.

Total CD45^+^ cells were increased about 50-fold in the lungs of hypereosinophilic compared to WT mice (Figure 3G). This was largely due to a massive increase in eosinophils, which accounted for almost half of all immune cells found in the lung (Figure 3H). In addition, neutrophils, monocytes, macrophages, CD4^+^ and CD8^+^ T cells were also increased in IL-5Tg lungs (Figure 3I-K and data not shown). Thus, hypereosinophilic mice develop extensive infiltration of the lungs that worsens with age and is about 50% eosinophilic. This finding mirrors the situation of HES patients where pulmonary eosinophil infiltration is found in over half of all patients and may be the presenting manifestation (*26*).

Pulmonary inflammation can change the lung architecture and increase pulmonary resistance. The resulting pulmonary hypertension increases the workload for the right ventricle and can eventually lead to right heart failure (*27*). To further investigate whether this process occurs in hypereosinophilic mice, we conducted echocardiography of the right ventricle and pulmonary valve on a cohort of older mice (age 30-41 weeks). To our surprise, we could not detect the typical signs of pulmonary hypertension. Pulmonary acceleration time was increased in IL-5Tg mice (Figure 3L). Pulmonary ejection time and the ratio of acceleration / ejection time were not significantly different between the two genotypes (Figure 3M, N). Moreover, mean pulmonary arterial pressure was decreased in IL-5Tg mice (Figure 3O). Thus, hypereosinophilic mice did not suffer from pulmonary hypertension. We conclude that while lung inflammation is very common in IL-5Tg mice and increases with age, it is probably not the cause of the cardiac pathologies that we observed.

### Eosinophil infiltration of the heart in IL-5Tg mice

To assess direct changes to the heart in hypereosinophilic mice we imaged histological sections of WT and IL-5Tg mice aged 30-41 weeks. While no gross abnormalities to the structure were apparent, hearts from hypereosinophilic mice showed a clear increase in immune cells in the cardiac muscle as compared to WT mice (Figure 4A, B). Infiltration appeared also more pronounced in these older mice as compared to young IL-5Tg mice (Figure 1N, O). We quantified the infiltrating CD45^+^ immune cells using flow cytometry. There was a 10-fold increase in total CD45^+^ cells in hypereosinophilic mice (Figure 4C). The vast majority (70%) of infiltrating cells were eosinophils, which were 100-fold more abundant in the hearts of hypereosinophilic mice compared to WT mice (Figure 4D). In contrast to the infiltrate in the lung which showed significant increases in multiple immune cell types, most other immune cell types were present at similar numbers in the hearts of IL-5Tg and WT mice (Figure 4E-H). The cardiac infiltrate in older hypereosinophilic mice is thus predominantly eosinophilic.

**Figure 4:**
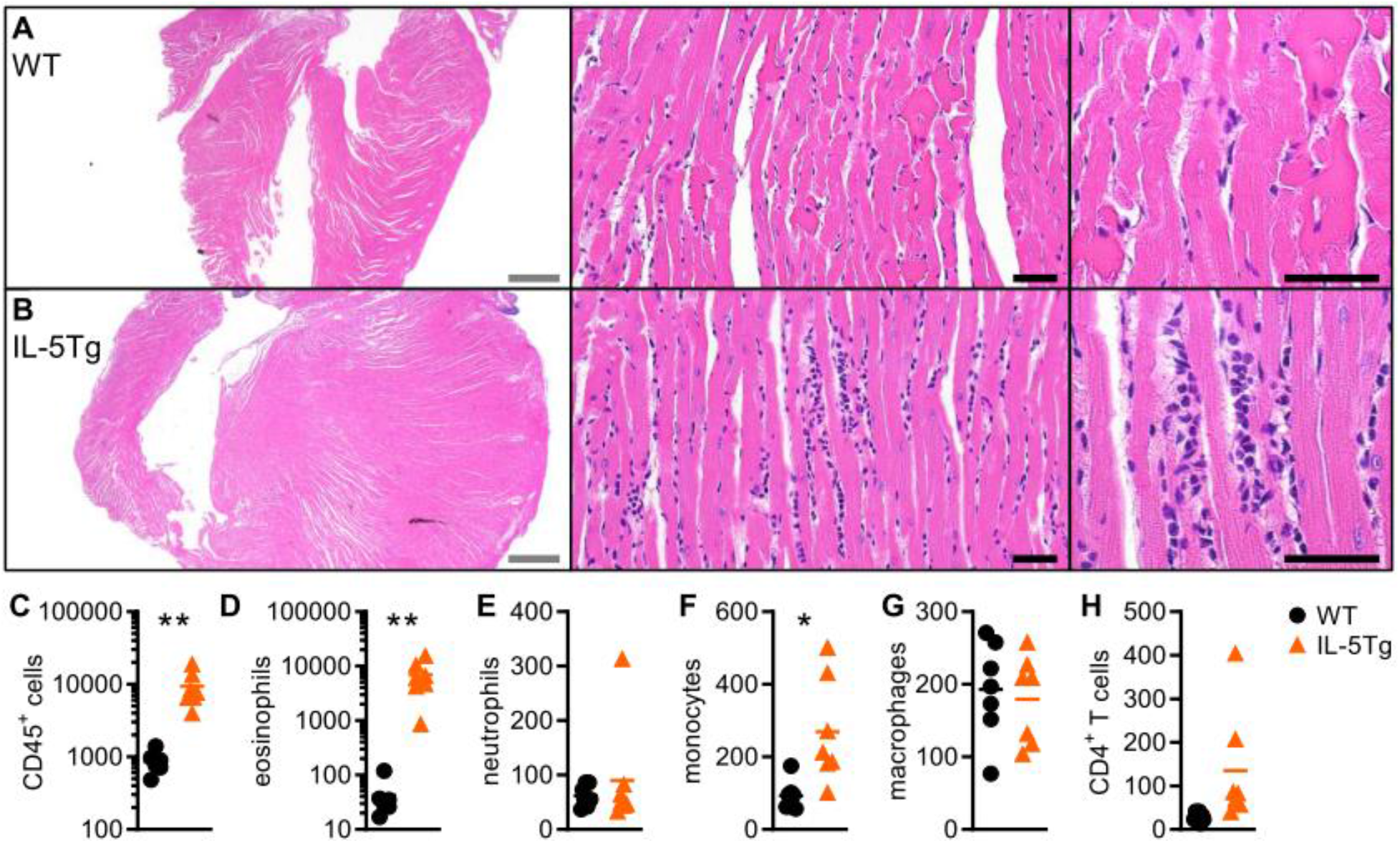
Older IL-5Tg mice develop eosinophilic infiltration of the heart. A-B) Representative histology images of A) WT and B) IL-5Tg hearts from mice aged 30-41 weeks. Grey scale bars: 1mm, black scale bars: 40µm. C-H) Infiltrating immune cells in the heart were quantified by flow cytometry in mice aged 30-41 weeks. Data are from 7 mice per genotype and were analyzed by t-test. * p<0.05, ** p<0.01.

### Cardiac eosinophils from IL-5Tg mice have an activated phenotype

We further characterized the infiltrating eosinophils in IL5Tg mice. In WT mice, eosinophils formed two populations, SiglecF-Lo and SiglecF-Hi that each accounted for about half of the eosinophils (Figure 5A). In IL-5Tg mice, eosinophils were predominantly SiglecF-Hi. IL-5Tg cardiac eosinophils had lower side scatter, which could result from increased degranulation (Figure 5B). CD11b expression was increased in the SiglecF-Hi compared to SiglecF-Lo cells of both genotypes (Figure 5C). Since IL-5Tg eosinophils were mostly comprised of the SiglecF-Hi subset they expressed overall higher levels of CD11b, a marker of eosinophil activation. CD45 signaling has been shown to limit cytokine expression in innate immune cells (*28*) and was reduced in IL-5Tg eosinophils. Compared to WT eosinophils, those from IL-5Tg hearts expressed lower levels of CD101 and CD62L, a marker of tissue-resident lung eosinophils that is reduced in inflammatory eosinophils (*29*) (Figure 5E, F). PDL1 expression, shown to increase after bacterial infection (*30*), was higher in IL-5Tg eosinophils (Figure 5G). Overall, this shows that cardiac eosinophils from IL-5Tg mice display an activated phenotype.

**Figure 5:**
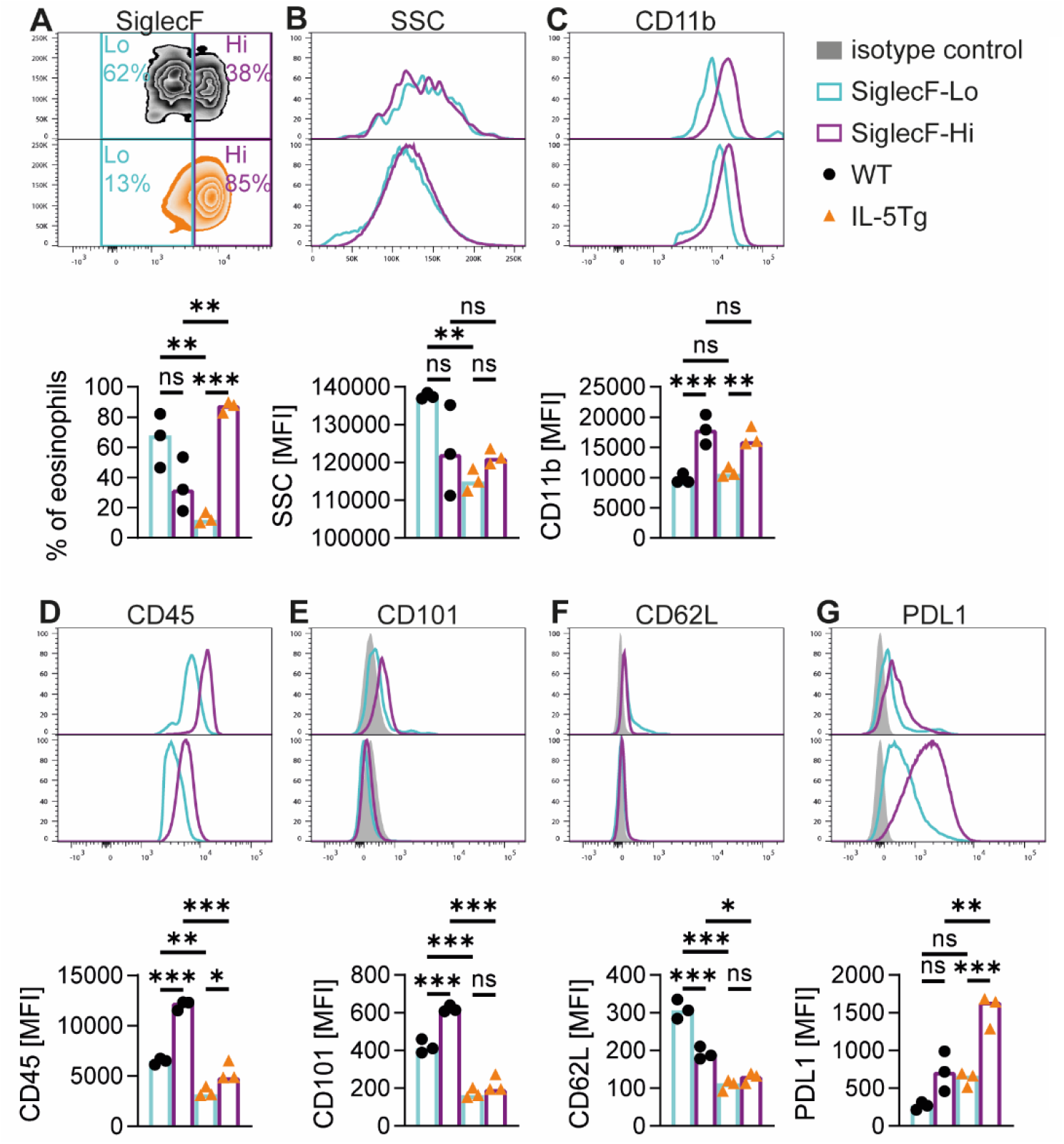
Cardiac eosinophil phenotype in IL-5Tg mice. Flow cytometry of eosinophils from the heart of 18 weeks old WT and IL-5Tg mice. Eosinophils were gated as live, CD45^+^CD11b^+^MHCII^-^ Ly6G^-^SiglecF^+^ cells. A) Frequency of SiglecF-Hi and SiglecF-Lo eosinophils in WT and IL-5Tg hearts. B-G) Mean fluorescent intensity (MFI) of selected markers in the SiglecF-Lo and SiglecF-Hi populations from WT and IL5Tg hearts. Data are from one experiment with n=3 mice per genotype. * p<0.05, ** p<0.01, *** p<0.001, oneway ANOVA with Sidak correction for multiple comparison.

### Gene expression profiling reveals an inflammatory signature in IL-5Tg hearts

We hypothesized that eosinophil infiltration will alter cardiac function on the molecular level. We performed gene expression analysis of left ventricular tissue from WT, IL-5Tg and IL-5TgΔdblGATA1 mice to identify underlying changes associated with impaired cardiac function. Samples clearly clustered by genotype in principle component analysis (PCA, Figure 6A). This revealed a major effect of IL-5 along PC1 with wildtype mice clustering away from IL-5Tg and IL-5TgΔdblGATA1 mice. The effect of eosinophils is apparent along PC2 with the longest distance between hypereosinophilic and eosinophil-deficient mice. Next, we examined differentially expressed genes in comparison to WT mice. Only 17 genes were commonly up-or downregulated in IL-5Tg and IL-5TgΔdblGATA1 mice, and 65 genes were uniquely differentially expressed in IL-5TgΔdblGATA1 mice (Figure 6B). The most dramatic changes occurred in IL-5Tg mice, where expression was changed in 172 genes, most of them being upregulated in comparison with WT mice (Data S1).

**Figure 6:**
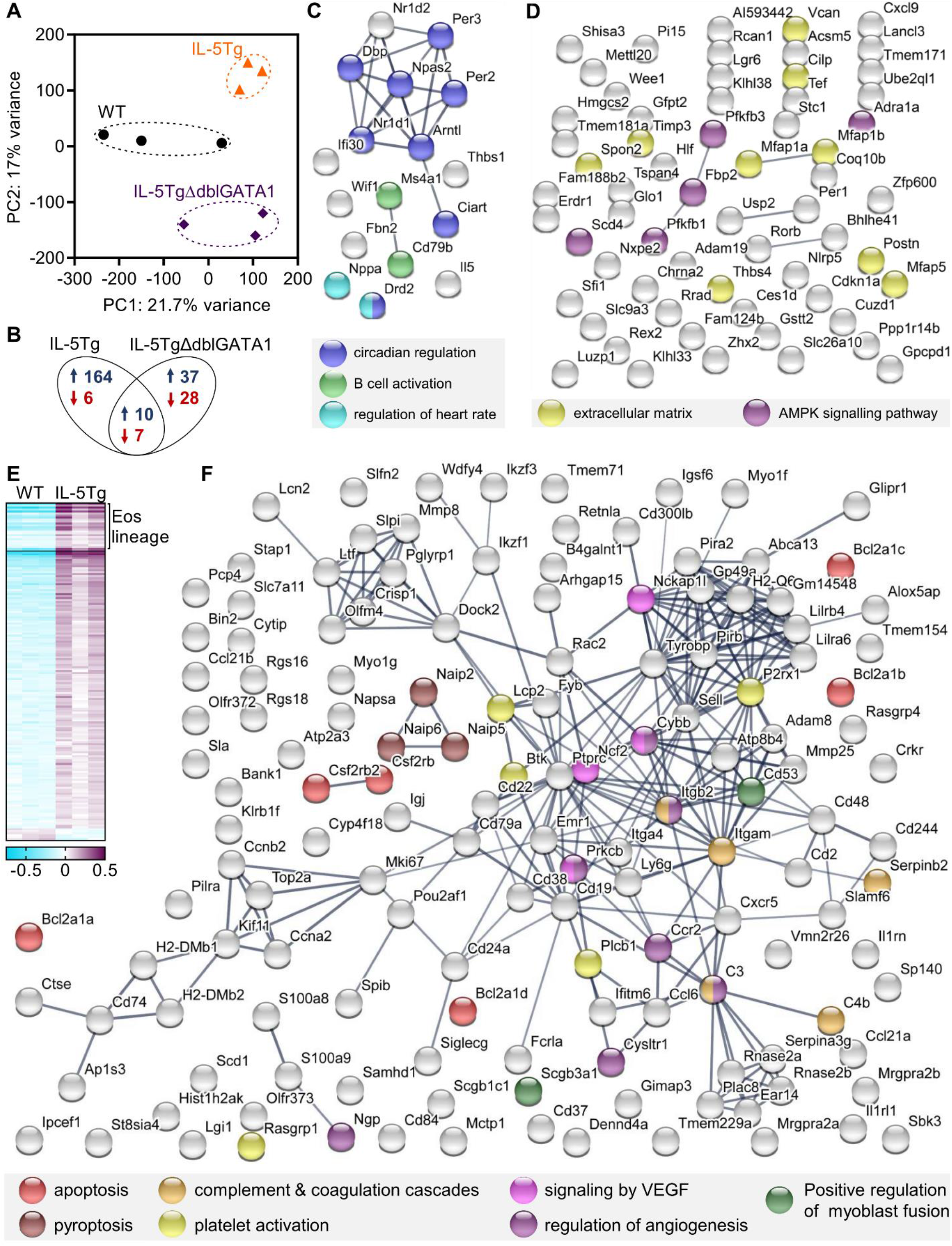
Hypereosinophilia leads to an inflammatory and tissue remodeling gene expression signature in the heart. Left ventricular tissue from mice aged 10-16 weeks was analyzed by microarray. A) Principal component analysis of cardiac gene expression data from different genotypes. B) Venn diagram of differentially expressed genes (p<0.05, |FC|≥1.4) in comparison to WT mice. C, D) Network analysis of protein-coding genes that are differentially expressed compared to WT mice (p<0.05, |FC|≥1.4) in C) both IL-5Tg and IL-5TgΔdblGATA1, D) only in IL-5TgΔdblGATA1. E) Differentially expressed genes (p<0.05, |FC|≥1.4) in IL-5Tg versus WT mice. Gene signature for eosinophils was generated using haemosphere database. F) Network analysis of protein-coding genes that are differentially expressed in IL-5Tg compared to WT mice, excluding genes that are specific to eosinophils. Network analysis was conducted using STRING database and proteins are color-coded for selected, significantly enriched gene ontology and KEGG pathway terms.

Network analysis of these differentially expressed genes revealed significantly more protein-protein interactions than would be expected (p=0.014 for IL-5TgΔdblGATA1; p<10^−16^ for IL5Tg and commonly regulated genes). Genes that were commonly up-or downregulated in both strains compared to WT mice are likely responsive to IL-5 but independent of eosinophils. As expected, this includes *Il5* itself (Figure 6C). It also includes genes associated with B cell activation, which are known to be increased in mice with high IL-5, and several circadian regulators. Interestingly, *natriuretic peptides A* (*Nppa*), a key regulator of cardiovascular homeostasis was downregulated in both IL-5Tg and IL-5TgΔdblGATA1 mice.

Genes that were differentially expressed only in IL-5TgΔdblGATA1 mice were enriched for factors associated with the extracellular matrix (Figure 6D). These include tissue remodeling factors *Timp3, Postn* and *Thbs4. Postn* (Periostin) is also known to be upregulated in activated cardiac fibroblasts (*31, 32*). Genes in the AMPK signaling pathway were also enriched in IL-5TgΔdblGATA1 mice. The alpha-adrenergic receptor *Adra1a* and several enzymes involved in glycolysis were upregulated, suggesting a possible increase in glycogen consumption.

We focused our attention on genes that were only differentially expressed in IL-5Tg mice. The large proportion of genes related to the immune system, all of which were upregulated in IL-5Tg compared to WT mice, stood out immediately (Supplemental Figure 3A). Many gene ontology terms related to immune processes were significantly enriched such as leukocyte migration, cytokine signaling, phagocytosis, lymphocyte activation and leukocyte differentiation, confirming an inflammatory environment in IL-5Tg hearts (Supplemental Figure 3B). As expected, eosinophil-specific genes (including *Epx, Prg2, Ccr3, Siglecf, Il5ra*) were upregulated in IL-5Tg mice (Supplemental Table 1). We also noticed several eosinophil-associated protein clusters, for example of eosinophil ribonucleases, of genes required for the synthesis of eoxins and leukotrienes, and those related to specific granules and respiratory burst (Supplemental Figure 3A). For example, the genes *Ncf1* and *Ncf2* are required for activation of the NADPH oxidase complex which produces superoxide. We also noted upregulation of *Ccl6*, which is produced by eosinophils and can cause an increase in reactive oxygen species in other cells (*33*). These eosinophil-specific clusters were clearly a result of the eosinophil infiltration into the heart tissue already identified by flow cytometry. Taken together, this suggests that the heart tissue in hypereosinophilic mice is constantly exposed to potentially damaging effects of eosinophils and other immune cells.

### Tissue damage and remodeling signatures in the hearts of hypereosinophilic mice

We also noted several gene ontology terms that suggest ongoing tissue damage and regeneration, such as apoptosis, platelet activation, regulation of angiogenesis and wound healing (Supplemental Figure 3). To gain a better understanding of the changes that happen to the heart tissue itself, we next excluded eosinophil-specific genes from the list of differentially expressed genes. Rather than manually selecting them, we used the haemosphere database to generate a list of eosinophil lineage genes. The resulting 157 genes are either specific to or expressed at much higher levels in eosinophils compared to other samples in the haemosphere database. Out of 172 genes with differential expression between IL-5Tg and WT hearts, 24 were eosinophil lineage genes (Figure 6E). We re-analyzed the remaining differentially expressed genes for protein networks and gene ontology enrichment. Immune system related genes remained significantly enriched. In addition, a clear signature of tissue damage and remodeling was now apparent. Remaining genes were enriched for those associated with apoptosis (*Bcl2* family) and pyroptosis (Nlrc4 inflammasome components) and the activation of complement and coagulation cascades (Figure 6F). *Alox15*, an eosinophil gene known to induce oxidation of phospholipids exposed on the outer plasma membrane for initiation of thrombi formation (*16*) was upregulated in IL-5Tg hearts (Supplemental table 1).

The second factor implicated in eosinophil-mediated coagulation initiation, tissue factor (gene *F3*) was detectable and significantly upregulated in IL-5Tg mice but did not reach our cutoff for fold enrichment. We also noted several extracellular matrix remodeling genes (*Mmp8, Mmp25, Adam8*), known to contribute to heart damage (*34-37*). Genes associated with VEGF signaling and angiogenesis were enriched, suggesting the growth of new blood vessels, a key feature of tissue remodeling. Proteins involved in cell proliferation (*Mki67, Ccna2, Ccnb2, Top2a, Kif11*) formed another cluster indicative of tissue repair and renewal. Lastly, myofibroblast fusion-related genes were enriched. This included *Cd53*, which is required for efficient formation of myofibers in regenerating muscle at the level of cell fusion (*38*). Inflammatory stress caused by eosinophils likely causes continuous damage to the heart muscle, which leads to an ongoing remodeling and repair process. We propose that this leads to the functional impairments in the cardiac conductive system and heart function that we observed in hypereosinophilic mice. This process could also explain the worsening of the phenotype with age, as damage accumulates and the heart fails to maintain its function.

### Cardiomyocytes from IL-5Tg mice exhibit impaired Ca^2+^ cycling and contractility

We hypothesized that the range of different cardiac pathologies that develop in hypereosinophilic mice may be caused by underlying mechanical abnormalities of the sarcomere and dysregulation in calcium cycling by cardiomyocytes. We isolated individual cardiomyocytes from WT and IL-5Tg mice, paced them (1Hz, 20V, 15ms), and measured contractility and Ca^2+^ transients. Cardiomyocytes from IL-5Tg mice exhibited significantly reduced contractility relative to WT controls, with reduced % shortening and faster relaxation kinetics (Figure 7A-D). The impairment in contractility was coupled with reduced Ca^2+^ release (Figure 7E-F), faster Ca^2+^ release kinetics (Figure 7G), and slower Ca^2+^ reuptake (Figure 7H). Our results suggest that dysregulation of contractility and Ca^2+^ reuptake in cardiomyocytes contributes to cardiac dysfunction at the whole organ level in hypereosinophilic mice.

**Figure 7.**
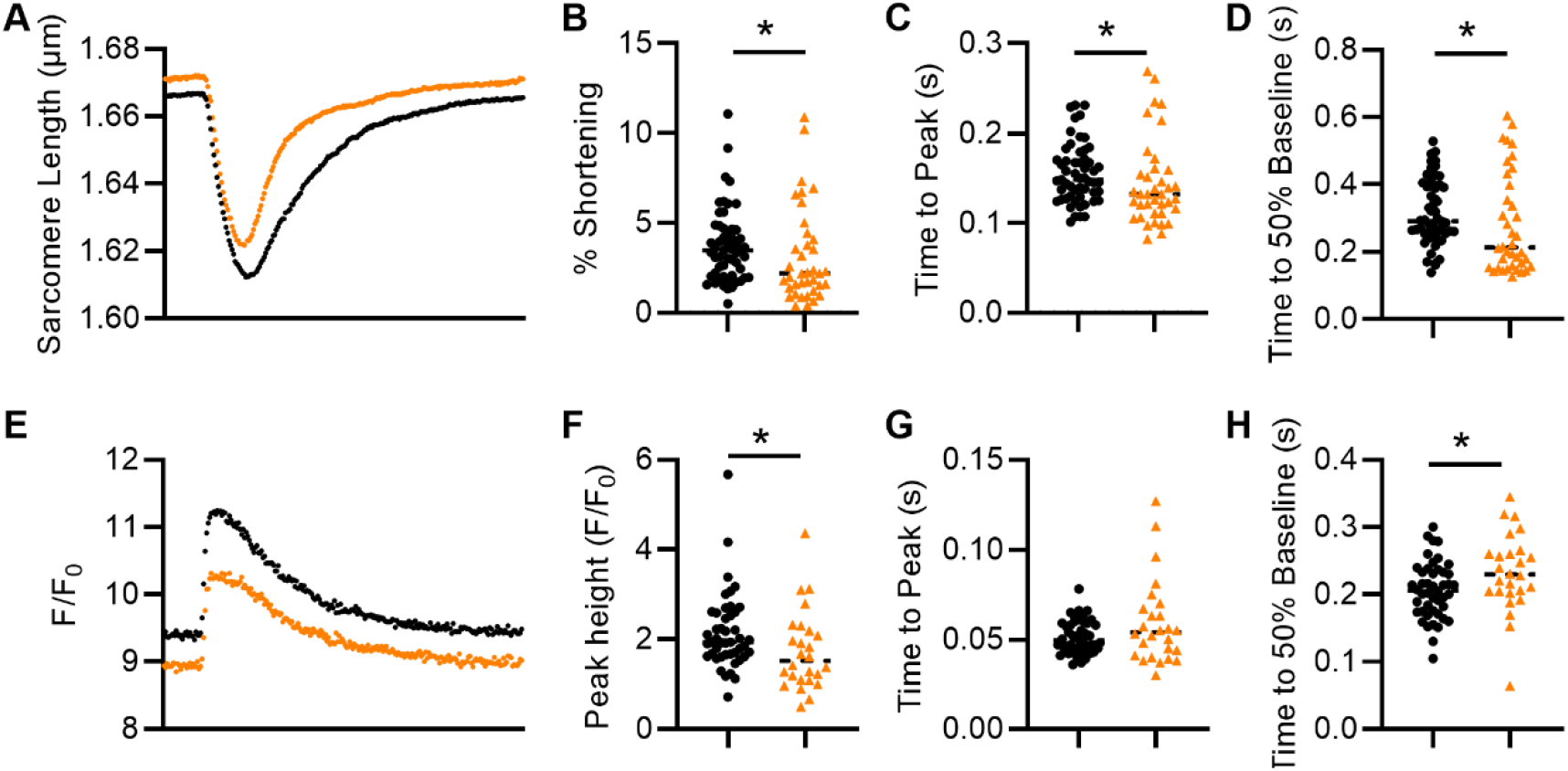
Cardiomyocytes from hypereosinophilic mice exhibit impaired contraction and relaxation kinetics. Sarcomere length shortening (A-D) and calcium transients (E-H) were measured in unloaded isolated cardiomyocytes. (A) Average trace of WT (black) and IL-5Tg (orange) cardiomyocyte sarcomere length shortening over 1 second. Quantification of % shortening (B), time to peak contraction (C), and time to 50% baseline (D). (E) Average traces of WT and IL-5Tg cardiomyocyte Ca^2+^ transients over 1 second. Quantification of maximal peak height of Ca^2+^ release (F), time to peak Ca^2+^ release (G), and time to 50% baseline for Ca^2+^ (H) show impaired Ca^2+^ handling. Each datapoint is from a single cardiomyocyte. 6-8 cardiomyocytes from n=3 mice were analyzed per group. Data are from 1 experiment. * p<0.05, ** p<0.01, Mann-Whitney test.

## Discussion

This study is the first characterization of the cardiac phenotype in hypereosinophilic mice. It comprises reduced left ventricular function that progressively worsens with age, increased heart weight, altered cardiomyocyte contractility and tissue infiltration by activated eosinophils. Gene expression signatures of inflammation, reactive oxygen burst, coagulation and the resulting tissue damage and remodeling reveal the processes behind this phenotype.

Echocardiography in young (6-10 weeks old) IL-5Tg mice showed a relatively uniform phenotype of decreased left ventricular ejection fraction, increased left ventricular diameters and thickened walls. When we repeated the analysis in older mice (20-30 weeks old), the phenotype was more variable. On average, hypereosinophilic mice still displayed reduced ejection fraction and increased LV diameters, but the individual variability was much higher. LV wall thickness was increased in some hypereosinophilic mice in comparison to WT controls, but decreased in others, suggesting that these mice can develop a range of pathologies as they age. It is also important to note that a few IL-5Tg mice did not show any abnormalities in echocardiography and had a heart weight that was within the range of their wildtype littermates. This highlights the heterogeneity of the phenotypic presentation and models HES as some patients with hypereosinophilia never develop cardiac pathologies (*6, 7*).

The reduction in LVEF at earlier ages is consistent with contractile impairment at the cardiomyocyte level in the IL-5Tg mice. Contractile kinetics were impaired in IL-5Tg cardiomyocytes during both contraction and relaxation. Corresponding impairment in Ca^2+^ transients suggest that eosinophil-mediated signaling impacts proteins that mediate Ca^2+^ release and reuptake. The prolongation of calcium reuptake during diastole is a pronounced cellular change often seen in heart failure in human and mouse models (*39, 40*). This is despite a reduction in maximal calcium release, as evidenced by the decrease in peak height by IL-5Tg ventricular myocytes. Disruption in calcium cycling can be attributed to dysregulation of one of the many calcium handling proteins of the sarcoplasmic reticulum (*41, 42*). Our data indicate that IL-5 overexpression and an increased population of eosinophils may negatively impact cardiomyocyte function leading to dysfunctional phenotypes seen in heart failure.

Our study offers clear mechanistic insight into how eosinophils drive cardiac pathologies. Gene expression data reveals signatures of tissue damage and inflammatory cell death through respiratory burst as well as specific eosinophil granules. These mechanisms have been proposed in the past as part of the pathogenesis in HES (*14*). Of particular interest is the complement and coagulation cascade. A recent study clearly demonstrated the mechanisms by which eosinophils promote coagulation (*16*). We find increased expression of both, Alox15, the enzyme responsible for generating the oxidized phospholipid surface, and of tissue factor in the hearts of hypereosinophilic mice. This activation of coagulation cascades relates to thrombotic events in HES patients (*5*) and possibly other eosinophilic diseases (*43*). Beyond eosinophilic diseases, COVD-19 infection and disease severity is clearly associated with eosinopenia in hospitalized patients (*44, 45*) and COVID-19 infections are marked by coagulopathy, in particular thrombotic microangiopathy (*46*). However, there is currently no evidence that eosinophils contribute directly to disease pathology or antiviral defense in this disease (*47*).

These gene expression data were generated in relatively young mice (11-15 weeks). At this point, LV function is already impaired but infiltration in the heart is still mild. It is possible that gene expression of older mice (30-40 weeks) would show even stronger gene signatures of tissue damage, remodeling and fibrosis. Nevertheless, this is the first gene expression analysis related to eosinophil-mediated heart damage. Bone marrow from HES patients has been sequenced (*48*), but to our knowledge there is no gene expression date from the heart in HES or other eosinophilic diseases.

In addition to gene signatures related to damage and inflammation we also found sings of tissue remodeling and repair. Several genes involved in the adverse remodeling of the extracellular matrix were upregulated in IL-5Tg hearts. The connection between eosinophils and fibrosis was made a long time ago (*49*) and is clearly important in eosinophil-mediated pathology in many organ systems (*24, 50-52*). Chronic inflammation and remodeling could also be directly linked to the pro-coagulative state generated by eosinophils (*53*). Additional gene expression signatures related to angiogenesis and VEGF signaling could be a direct consequence of eosinophils, which can produce VEGF and other pro-angiogenic factors (*54*). Angiogenesis forms part of the broader tissue regeneration and repair process that seems to be ongoing in the hearts of hypereosinophilic mice.

Recently, it has been shown that eosinophil recruitment to the heart is beneficial during myocardial infarction (*55-57*). This seems at first contradictory to our findings presented here but it is likely that different mechanisms are at play. Myocardial infarction results in substantial tissue damage and an appropriate level of tissue remodeling is needed. Eosinophils have been shown to promote tissue regeneration in other organs (*58, 59*) and it is possible that they fulfill a similar role in the heart post myocardial infarction. In these studies, wildtype mice were compared to eosinophil-deficient ddblGATA1 mice or supplemented with IL-5 and eosinophils played an anti-inflammatory and immune regulatory role, limiting cardiomyocyte death and promoting alternative macrophage activation and angiogenesis. Interestingly, we found gene expression signatures of VEGF signaling and regulation of angiogenesis in our dataset. However, as we showed here, eosinophils from hypereosinophilic mice have an activated phenotype which may be the reason for their negative effects on cardiomyocytes. It is possible that the effect of eosinophils is dependent on their activation status and microenvironment as well as on the context, such as an acute versus chronic model. Together these studies and our findings show that the role of eosinophils in the heart is context-dependent and that eosinophils may have both beneficial and detrimental functions.

The IL-5Tg mouse has its limits as a model for HES as it does not replicate the ‘classic’ cardiac presentation of HES with endomyocardial thrombi and fibrosis. While we did find relevant gene signatures, especially of coagulation, hypereosinophilic mice did not display the characteristic intraventricular thrombi or fibrosis. It is possible that these develop in older mice beyond 41 weeks of age, which was the limit of this study. The pulmonary infiltration in hypereosinophilic mice ranged from dispersed, interstitial eosinophils to dense nodules of mixed, predominantly eosinophilic infiltrate. This shows clear similarities to patients. A quarter of HES patients has pulmonary involvement, which is characterized by eosinophilic infiltrates that can form nodules and vascular proliferation (*60*). Given the dramatic lung infiltration, we were surprised not to find any signs of pulmonary hypertension by echocardiography. This suggests that the cardiac pathologies observed are due to heart inflammation directly and our subsequent analysis of cardiomycotye contraction kinetics highlights the causal processes. Nevertheless, we do not want to exclude the possibility that the widespread tissue infiltration that is known to occur in aging IL-5Tg mice also affects their cardiovascular system. In addition to lung infiltration, IL-5Tg mice display dramatic splenomegaly and have enlarged livers (*25*). These organs most likely have altered vasculature, although this has not been studied directly. Any such changes are prone to increase cardiac workload or resistance and could over time contribute to the observed cardiomegaly.

We should note that this study does not alter the conclusions drawn from our earlier publication on experimental autoimmune myocarditis in IL-5Tg mice (*61*). Experimental autoimmune myocarditis drives an antigen-mediated, T cell-driven inflammation that peaks after 2 weeks and leads to chronic remodeling after 6 weeks (*62*). In our previous paper, mice were immunized at 6-8 weeks and analyzed by 12-14 weeks. The spontaneous, progressive disease of IL-5Tg mice that we describe here develops on a much slower time scale. In addition, we had normalized to baseline heart function before immunization, thereby adjusting for the differences between WT and IL-5Tg mice we describe here in detail. The dilated cardiomyopathy that develops in IL-5Tg mice after induction of experimental autoimmune myocarditis is also much more uniform than the varied phenotypes that develop spontaneously in aging IL-5Tg mice (*61*).

Recently, Prows et.al. published a different mouse model of hypereosinophilia and heart disease. This spontaneous mouse mutant displays hypereosinophilia, tissue infiltration and eosinophilic myocarditis that results in sudden, premature death (*63*). As such, it is a more sudden, severe model compared to the slow, progressive disease we observe in IL-5Tg mice. The advantages of the IL-5Tg model is that it is genetically defined and well characterized in multiple studies (*16, 25, 33, 61*). Because of the known genetic alterations (CD3δ-driven IL-5 expression) it is possible to isolate the effect of eosinophils as we did here by using IL-5TgΔdblGATA1 mice. Importantly, both studies highlight the crucial role of eosinophils in driving cardiac pathology.

In summary, we have characterized in detail the cardiac pathologies that develop in hypereosinophilic IL-5Tg mice including their progressive impairment of left ventricular function, abnormalities in cardiomyocyte contractility and the eosinophil infiltrate in the heart. Underlying changes in gene expression highlight the process of ongoing tissue damage and resulting remodeling that is likely insufficient to maintain cardiac function. This study is directly relevant to the disease pathogenesis in HES and beyond that to other eosinophilic diseases like eosinophilic myocarditis, eosinophilic granulomatosis with polyangiitis or bullous pemphigoid as well as broader cardiovascular disease, where eosinophils are associated with occurrence of thrombotic events (*16*) and atrial fibrillation (*64*).

## Material and Methods

### Mice

IL-5Tg mice (line NJ.1638, (*25*)) were kindly provided on the BALB/c background by Drs. James and Nancy Lee, Mayo Clinic, Scottsdale and maintained by breeding IL-5Tg with BALB/cJ wildtype mice. BALB/cJ wildtype (WT, Jackson Laboratories Stock #651) and ΔdblGATA1 (*65*), #5653), mice were purchased from Jackson Laboratories. All mice were on the BALB/c background. Mice were housed in specific pathogen-free animal facilities at the Johns Hopkins University. Experiments were conducted on age-matched male and female mice in compliance with the Animal Welfare Act and the principles set forth in the Guide for the Care and Use of Laboratory Animals. All methods and protocols were approved by the Animal Care and Use Committee of Johns Hopkins University.

### Histology and light microscopy

Heart and lung tissue was fixed in SafeFix solution (Thermo Fisher Scientific), embedded, and cut into 5-µm serial sections prior to hematoxylin and eosin staining. Images were acquired with an Olympus DP72 camera on an Olympus BX43 microscope using cellSens Standard version 1.4.1 (Olympus). Quantification of myocyte cross-sectional area was performed as previously described (*66*). In short, following deparaffinizing and citrate-based heat-mediated antigen retrieval, slides were incubated with Alexa Fluor 647-conjugated wheat germ agglutinin (W32466, Invitrogen) at 4°C overnight and mounted using Prolong Gold mounting medium (P36934, Thermofisher). Image acquisition was performed on an EVOS epifluorescence microscope (Life Technologies). The cross-sectional area of 2061-7603 cells from 5-9 areas per mouse heart was measured using an automated algorithm with Image J and averaged.

### Echocardiography

For echocardiography, conscious, depilated mice were held in a supine position. Transthoracic echocardiography was performed using the Vevo2100 or Acuson Sequoia C256 ultrasonic imaging systems (Siemens) with a 13 MHz transducer. The heart was imaged in the parasternal short axis view in the two-dimensional mode. An M-mode cursor was positioned perpendicular to the interventricular septum and the left ventricular posterior wall at the level of the papillary muscles. The left ventricular (LV) end-diastolic diameter (LVEDD), and LV end-systolic diameter (LVESD), interventricular septal (IVS) wall thickness at end-diastole, and LV posterior wall thickness at end-diastole were measured three times for each mouse from a frozen M-mode tracing and averaged using Vevo LAB 5.6.1 software. Ejection fraction (EF), relative wall thickness (RWT), and LV mass were calculated from these parameters as previously described (*67*).

### Flow cytometry

For flow cytometry analysis, mouse hearts and lungs were perfused for 3 min with HBSS containing no Ca^2+^ and Mg^2+^. Hearts were digested for 30 min at 37°C in gentleMACS C Tubes (Miltenyi Biotec) with 800 U/ml Collagenase II and 100 U/ml DNase I (Worthington Biochemical Corporation). Lungs were digested with 100 U/ml Collagenase II and 50 U/ml DNase I under same condition as hearts. To exclude dead cells, LIVE/DEAD staining was done according to the manufacturer’s instructions (Thermo Fisher Scientific). Cells were blocked with anti-CD16/CD32 (eBioscience) and were stained with fluorochrome-conjugated monoclonal antibodies (eBioscience, BD, and BioLegend). For absolute quantification, viable cells were counted CountBright beads (Thermo Fisher Scientific). Samples were acquired on an LSR II cytometer running FACSDiva 6 software (BD). Data were analyzed with FlowJo 10 (Tree Star).

### Mouse ventricular cardiomyocyte isolation

Adult IL-5Tg mice and litter mate control mice were anesthetized with 4% isoflurane, in accordance to approved JH IACUC protocols. Hearts were excised and cannulated via the aorta for retrograde perfusion. Hearts were perfused for 3 mins at 37°C in a buffer containing NaCl (120 mM), KCl (5.4 mM), MgSO_4_ (1.4 mM), NaH_2_PO_4_ (12 mM), NaHCO_3_ (20 mM), 2,3-butadiene monoxime (10 mM), taurine (5 mM), and glucose (5.6 mM) as previously described (*68-71*). Next, hearts were digested in a buffer containing Collagenase, Type II (0.25 mg/ml, Worthington) for 10-15 mins. The ventricles were removed and manually minced and pipetted before being filtered through a 200 µm mesh. The cardiomyocytes were resuspended in 1.8 mM Ca^2+^-Tyrode’s solution containing NaCl (140 mM), KCl (4 mM), MgCl_2_ (1 mM), glucose (10 mM), and HEPES (5 mM), pH 7.4, at room temperature and reintroduced to extracellular Ca^2+^ in a stepwise manner up to 1 mM calcium.

### Measurement of ventricular cardiomyocyte contraction and calcium kinetics

To perform functional analysis of cardiomyocytes, isolated single cardiomyocytes were sorted onto laminin-coated coverslips. Coverslips were then transferred to an inverted microscope (Nikon, TE2000) to measure sarcomere shortening and Ca2^+^ transient changes. Cells were loaded with leak-resistant Ca^2+^ indicator Fura-2 AM (Abcam) and measured using the IonOptix imaging system and software (IonOptix, Westwood, MA). Cells were paced at 1 Hz, at room temperature in Tyrode’s containing 1 mM Ca2^+^ and 0.01% DMSO, and both sarcomere length shortening and Ca^2+^ transients were assessed simultaneously.

### Microarray

Left ventricular tissue from naïve 11-15 weeks-old WT, IL-5Tg and IL-5TgΔdblGATA1 mice was homogenized in Trizol and total RNA isolated according to the manufacturer’s protocol. RNA transcript levels were quantified by microarray on the Affymetrix Mouse Transcriptome Array 1.0 GeneChip array. Generation of cDNA, second strand synthesis, and labeling used the Affymetrix Ambion WT Expression Kit (Affymetrix) according to the manufacturer’s protocol. These arrays were washed and stained using the GeneChip Fluidics station 450, labeled cDNA probes hybridized, and the arrays scanned using the GeneChip Scanner 3000 7G and default parameters (all Affymetrix). The Affymetrix Expression Console generated raw data as CEL files which were imported into the Partek Genomics Suite v6.6 (Partek Inc.) for analysis. Data were extracted using the MTA-1_0.na34.1.mm10.transcript annotation, quantile normalized with the RMA (Robust Multi-Array Average) algorithm, and log2 transformed, yielding data for 25,094 gene-annotated transcripts, whose annotation was subsequently updated to MGI/NCBI nomenclature. After data quality control to exclude possible outliers, the samples’ gene expression was independently compared across the three biologic classes using a two tailed one way ANOVA, which provided relative levels of gene expression and statistical significance, as fold change and p-value respectively. Protein-coding genes with p values <0.05 and fold changes >1.4 were considered differentially expressed and further analyzed. The reported microarray data are MIAME compliant and are accessible in the NCBI Gene Expression Ominbus under accession number GSE185300.

### STRING analysis

Lists of differentially expressed genes were uploaded to STRING database (www.string-db.org), version 11.0 and organism set to Mus musculus. Only high confidence (>0.7) interactions based on all available interaction sources were considered for the generation of networks. Line thickness indicates the strength of data support (thick: very high confidence (≥0.9), thin: high confidence (≥0.7)). For gene ontology and pathway analysis, the whole genome was assumed as the statistical background.

#### Generation of eosinophil gene signature using haemosphere database

We used the ‘high expression search’ on the haemosphere database (www.haemosphere.org) to generate a list of genes with unique or high expression in the eosinophil lineage in comparison to remaining cell types in the ‘Haemopedia-Mouse-RNASeq’ dataset. Results were used to exclude eosinophil-specific genes from our list of differentially expressed genes in IL-5Tg versus WT mice.

### Statistics

Two groups with normally distributed data were analyzed using student’s t-test. Mann-Whitney test was used for nonparametric data and data that were not normally distributed. Multiple group analysis was performed by one-way ANOVA followed by Tukey’s multiple comparisons test. Comparisons within each age group for outcomes recorded in multiple age groups were performed by one-way ANOVA followed by Holm-Sidak correction for multiple comparison. Fisher’s exact test was used to determine significant differences in the proportion of mice displaying pathologic features. Linear regression analysis was used to analyze correlation between parameters. Calculations were performed in Prism 9.2.0 (GraphPad Software Inc.). P values <0.05 were considered statistically significant and are denoted by asterisk: *p<0.05, **p<0.01, ***p<0.001.

## Author Contributions

Conceptualization: NLD, DC

Methodology: NLD, MKW, TW, WB-B, MVT, BLL, CL, DB, NW

Investigation: NLD, MKW, TW, WB-B, BLL Visualization: NLD, MKW, TW

Supervision: DC, DK

Formal analysis: CT Writing—original draft: NLD

Writing—review & editing: NLD, MKW, TW, BLL, DC

Funding acquisition: DC

## Data and materials availability

Microarray data will be available on GEO under accession number GSE185300.

## Supplementary Materials

Fig S1-S3

Movie S1-S2

Data S1

