## Supplemental Figures for "Hypereosinophilia causes progressive cardiac pathologies in mice"

**This PDF file includes:**

Figs. S1 to S3

**Other Supplementary Materials for this manuscript include the following:**

Movies S1 to S2
Data S1

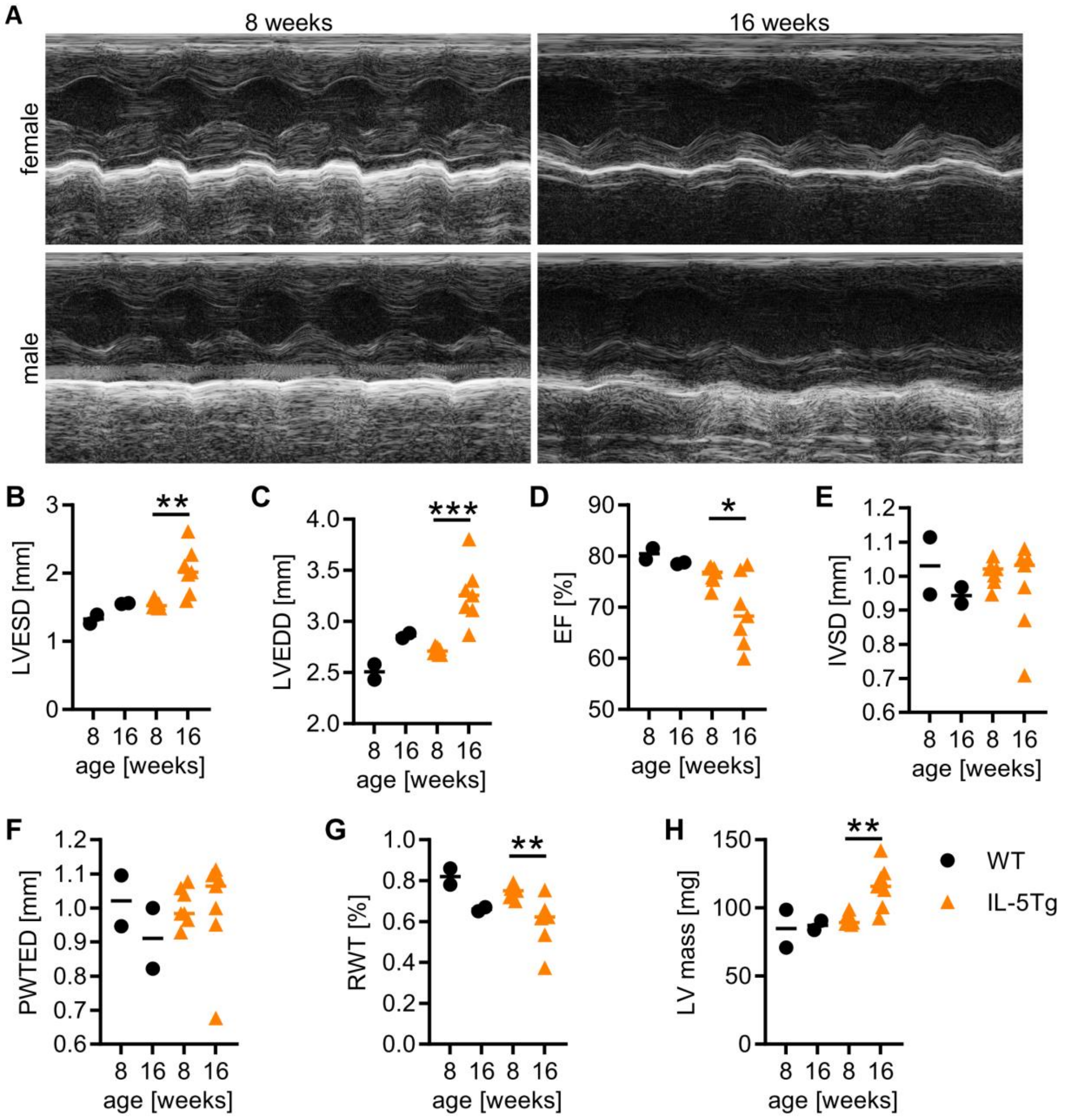


**Fig. S1. Age exacerbates cardiac pathology in hypereosinophilic mice**

A) Representative M-mode images from IL-5Tg mice at 8 and 16 weeks of age. B-H) Serial echocardiography was performed on n=7 IL-5Tg mice at 8 and 16 weeks of age and n=2 WT mice were included for reference. IL-5Tg mice were compared by t-test at 8 versus 16 weeks of age.

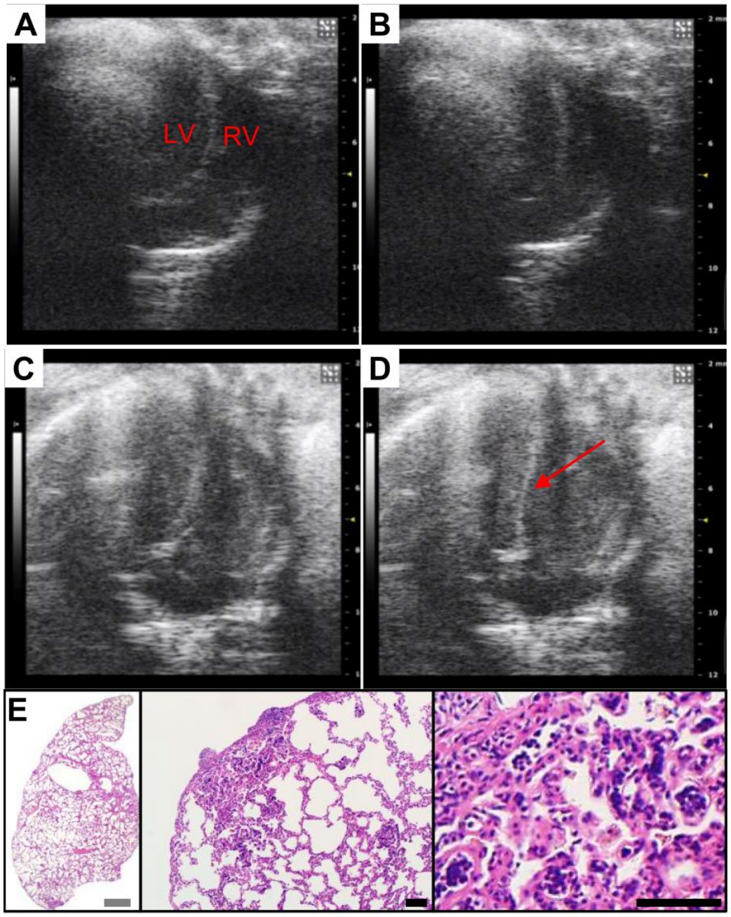


**Fig. S2. Right ventricular dilation and lung pathology in an old Il-5Tg mouse**.
A-D) Echocardiography 4-chamber view of a 46-weeks-old WT (A-B) and a 45-weeks-old IL-5Tg mouse (C-D). E) H&E-stained lung section of the IL-5Tg mouse. Grey scale bar: 1mm, black scale bar: 40µm

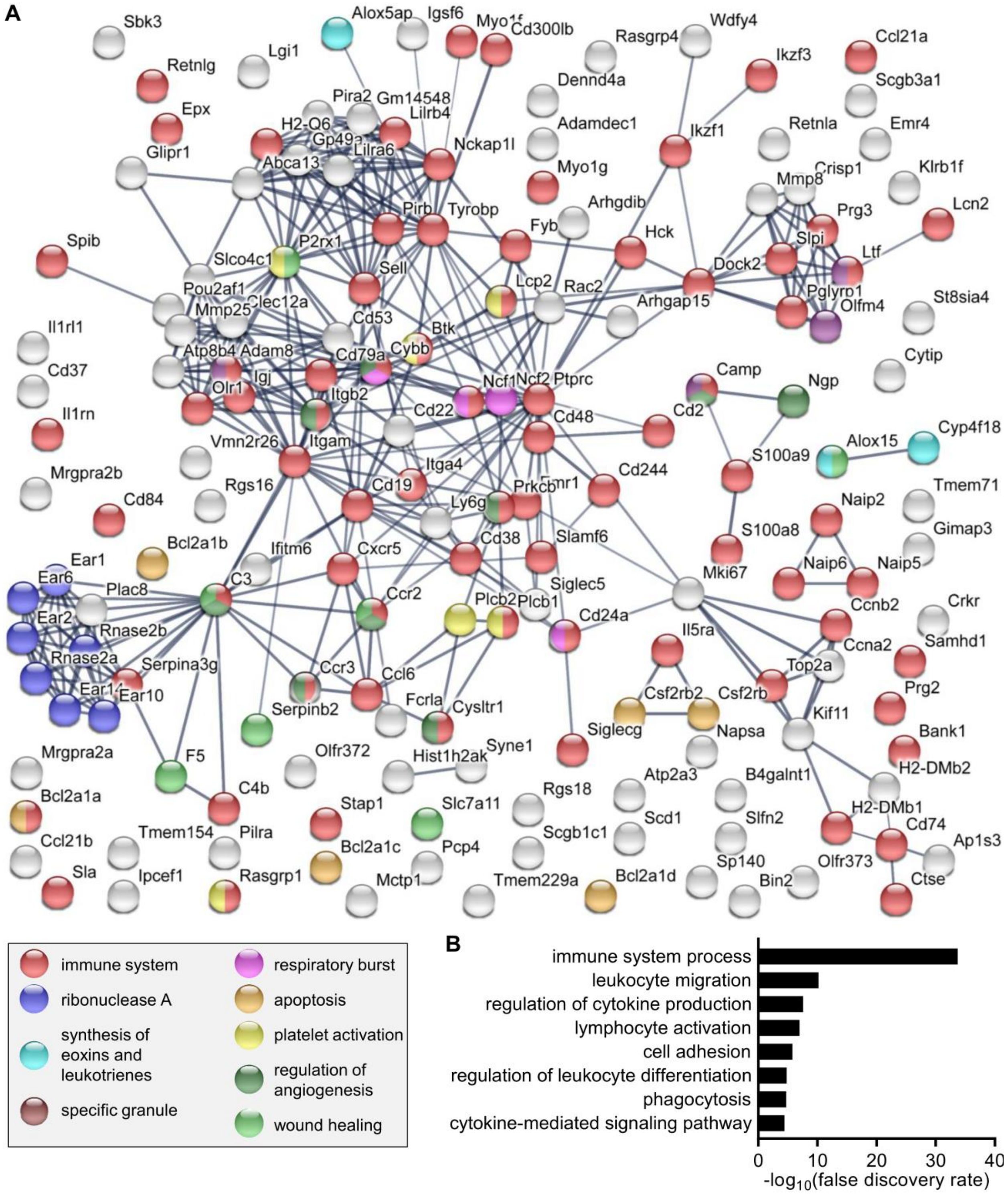


**Fig.S3. Signatures of inflammation and tissue damage in IL-5Tg hearts.**
Left ventricular tissue from mice aged 10-16 weeks was analyzed by microarray. Network analysis of protein-coding genes that are differentially expressed in IL-5Tg compared to WT mice ($p<0.05$, $|FC|\geq 1.4$). Analysis was conducted using STRING database. A) Proteins are color-coded for selected, significantly enriched gene ontology and KEGG pathway terms. B) Selected gene ontology terms from the analysis in A.

**Movie S1.**

Echocardiography, 4-chamber view of WT mouse aged 49 weeks.

**Movie S2.**

Echocardiography, 4-chamber view of IL-5Tg mouse aged 46 weeks.

**Data S1.**

Data table of differentially expressed genes and Haemosphere eosinophil lineage genes.
